# The microbial soy metabolite S-equol mitigates cadmium neurotoxicity in mice

**DOI:** 10.64898/2026.10.01.756095

**Authors:** Youjun Suh, Hao Wang, Lingjun Li, Haiwei Gu, Johannes Johnson-Martinez, Andrew T. Magis, Sean M. Gibbons, Julia Yue Cui

## Abstract

Environmental exposures are major but understudied determinants of chronic disease. Cadmium is a pervasive toxic metal encountered through diet and tobacco smoke that has been associated with cognitive impairment and Alzheimer’s disease mortality, yet strategies to mitigate its neurotoxicity remain limited. S-equol is a gut microbial metabolite of soy daidzein that crosses the blood brain barrier and preferentially activates estrogen receptor beta. Here, we show that cadmium disrupts an axis linking the equol producing gut bacterium *Adlercreutzia equolifaciens* to circulating S-equol and hippocampal estrogen receptor expression in mice, that S-equol attenuates cadmium-induced cytotoxicity and restores neuronal, glutamatergic and synaptic transcriptional programs in mouse neural progenitor cells, and that circulating S-equol in older humans is lower in apolipoprotein E4 carriers and covaries with body mass index, identifying S-equol as a microbiome-derived, host-modulated candidate countermeasure against cadmium neurotoxicity.

## INTRODUCTION

Alzheimer’s disease (AD) is a multifactorial neurodegenerative disorder whose risk is shaped by interactions among genetic susceptibility, aging, and environmental exposures. In the present study, cadmium (Cd) was selected as a representative environmental risk factor because of its widespread human exposure, established neurotoxicity, and epidemiological association with AD-related mortality. In U.S. adults aged ≥60 years, blood Cd concentrations >0.6 μg/L were associated with a 3.83-fold higher hazard of AD mortality compared with concentrations ≤0.3 μg/L^1^. Human exposure to Cd occurs predominantly through food in nonsmokers and through both dietary sources and tobacco smoke in smokers^2^, with cereals and bread, leafy vegetables, potatoes, legumes and nuts, and root and stem vegetables representing major dietary contributors in the U.S. population^3^. Cd neurotoxicity is mediated through multiple mechanisms, including disruption of calcium and zinc homeostasis, impairment of blood–brain barrier integrity, mitochondrial dysfunction, oxidative stress, neuroinflammation, altered neurotransmission, and neuronal injury^4, 5, 6^. Consistent with these mechanisms, we showed that experimental Cd exposure in mice impairs learning and memory and suppresses adult hippocampal neurogenesis^7, 8^. Increasing evidence further implicates the gut microbiome as an important mediator of Cd toxicity. Our previous work demonstrated that oral Cd exposure disrupts gut microbial composition and metabolic function in humanized *APOE*3/E4 transgenic mice, with pronounced perturbation of the gut-liver axis in *APOE*4 males^9^. The human *APOE*4 allele is the strongest known genetic risk factor for AD^10, 11^. More recently, we showed that Cd-induced gut dysbiosis preceded the development of hippocampus-dependent learning and memory deficits and was accompanied by impaired intestinal barrier-related gene expression, inflammatory activation, altered neuroactive microbial metabolites, and hippocampal transcriptional changes related to cognition and neuroinflammation^12^. Collectively, these findings support a model in which Cd may influence neurological vulnerability not only through direct neurotoxic effects, but also through disruption of the microbiome-gut-brain axis, raising the possibility that microbiome-derived metabolites represent modifiable mediators of Cd-associated neurotoxicity.

S-equol is a bioactive gut microbial metabolite generated through the stereospecific reduction of daidzein, one of the principal isoflavones in soy, with human intestinal bacteria producing almost exclusively the S-enantiomer^13^. Several equol-producing bacteria have been isolated directly from the human intestine, including *Adlercreutzia equolifaciens*, *Slackia isoflavoniconvertens*, *Slackia equolifaciens*, and *Eggerthella sp. YY7918*^14, 15, 16, 17^. Among soy isoflavones and their metabolites, S-equol is considered one of the most biologically potent and displays particularly strong estrogenic activity, preferentially activating estrogen receptor-β (ERβ) at nanomolar concentrations; its affinity for ERβ is substantially greater than that of its precursor daidzein^13, 18, 19^. This ERβ-predominant activity, together with potent antioxidant, anti-inflammatory and neuroprotective properties, has generated considerable interest in S-equol as a microbiome-derived modulator of brain health^18, 19, 20^. S-equol crosses the blood-brain barrier and protects neuronal cells against inflammatory and neurotoxic insults, and S-equol-containing ERβ-selective phytoSERM interventions enhance mitochondrial respiration and bioenergetic enzyme activity and reduce oxidative damage in experimental brain models^20, 21^; S-equol also protects against Aβ-induced neuronal injury and ERβ-dependent synaptic loss in experimental models^22, 23^. Following absorption, however, S-equol undergoes extensive phase-II metabolism, predominantly glucuronidation by UDP-glucuronosyltransferases (UGTs), with human liver microsomes displaying greater intrinsic glucuronidation capacity than intestinal microsomes; consequently, <1% of circulating S-equol is present in its unconjugated form and a large fraction of an oral dose is ultimately recovered in urine, consistent with efficient systemic elimination^24, 25, 26, 27^. Importantly, despite its estrogenic activity, available human evidence does not indicate an increased cancer risk associated with S-equol: clinical supplementation studies have not identified concerning effects on breast or endometrial endpoints^28, 29^, and observational studies have instead associated higher endogenous equol exposure with lower risks of breast and prostate cancer^30, 31^.

Epidemiological and intervention studies examining soy consumption and cognitive decline or dementia have yielded inconsistent results: higher midlife tofu consumption was associated with poorer late-life cognition and greater brain atrophy in Japanese-American men^32^, whereas a large prospective Japanese cohort found no association between total soy or isoflavone intake and disabling dementia overall, with a possible inverse association for natto in women^33^, and long-term soy-isoflavone supplementation failed to improve global cognition in a randomized trial of postmenopausal women^34^. One potential explanation for these discordant findings is that dietary soy exposure does not necessarily translate into S-equol exposure: approximately 40-70% of East Asians, including ∼40-50% of contemporary Japanese populations, are S-equol producers, compared with only ∼20-30% of Western populations^19, 35^, with U.S. studies reporting producer frequencies of approximately 27-30%^36, 37^. Equol-producing capacity is fundamentally a microbial phenotype and can be influenced by diet: vegetarians have been reported to exhibit substantially higher equol-producer frequencies than nonvegetarians (59% versus 25%)^38^, while daidzein intake in a large Japanese cohort was associated with equol-producing status and increased abundance of equol-associated intestinal bacteria^39^. Thus, long-standing population differences in dietary exposure and microbiome structure may contribute to geographic variation in S-equol-producing capacity, providing a plausible explanation for why studies based solely on soy-product intake may yield heterogeneous neurological outcomes across populations. Consistent with this concept, a randomized trial of soy isoflavones in patients with AD found no overall cognitive benefit, yet exploratory analyses linked higher plasma equol concentrations with better verbal fluency and speeded dexterity^40^, while among cognitively normal older Japanese adults, equol-producing status was inversely associated with cerebral white-matter lesion burden, although not with amyloid-β positivity^41^. Moreover, a small pilot study in women with AD provided preliminary evidence of mitochondrial target engagement following S-equol administration, with 11 of 15 participants showing the prespecified positive platelet cytochrome-c oxidase response, although the study was not designed to establish cognitive outcomes^42^; a larger 24-month randomized trial involving 369 older adults is now directly evaluating S-equol effects on brain white-matter lesions and cognition^43^.

Despite these emerging links between S-equol, mitochondrial function and brain health, whether environmental neurotoxicants modify microbiome-dependent S-equol availability, and whether S-equol in turn modifies their neurotoxicity, remain largely unexplored. We therefore investigated whether Cd perturbs microbiome-associated S-equol availability and whether restoration of S-equol can mitigate Cd-induced neural toxicity, thereby testing S-equol as a potential microbiome-derived mediator of environmental susceptibility in the gut-brain axis.

Here, we show that cadmium perturbs microbiome-associated S-equol availability and that restoring S-equol mitigates cadmium induced neural injury. Oral cadmium exposure reduces *Adlercreutzia equolifaciens* abundance and hippocampal estrogen receptor expression in mice; S-equol attenuates cadmium cytotoxicity and restores neuronal, glutamatergic and synaptic transcriptional programs in neural progenitor cells; and in older adults, circulating S-equol is lower in *APOE4* carriers and covaries with body mass index. Together these results identify S-equol as a candidate microbiome-derived mediator of environmental susceptibility in the gut-brain axis.

## RESULTS

### Cd disrupts the microbiome-S-equol - estrogen receptor axis

To determine to what extent Cd exposure alters the intestinal source and circulating abundance of S-equol, we examined the major equol-producing intestinal microbe *Adlercreutzia equolifaciens* by shallow shotgun metagenomic sequencing (MGS), serum S-equol (LC-MS), and hippocampal estrogen receptor expression (RNA-Seq) in mice following 9-weeks of 3 mg/L CdCl₂ in drinking for 9 weeks. To note, we previously reported that this dosing regimen resulted in blood Cd levels similar to smokers^44, 45^. Cd exposure significantly decreased the centered-log ratio (CLR) abundance of *A. equolifaciens* compared with controls (Fig. 2A). To note, *A. equolifaciens* is also a well-characterized S-equol-producing bacterium in the human intestine^14, 46^. Consistent with this microbial change, circulating S-equol showed a downward trend in Cd-exposed mice, although the difference did not reach statistical significance (Fig. 2B). To note, unlike humans, in whom only a subset of individuals harbors gut microbiota capable of producing equol, all conventionally raised laboratory mice are generally efficient equol producers^27, 47, 48^. Thus, the observed decreasing trend in circulating S-equol is unlikely to reflect an absence of equol-producing capacity and instead supports a Cd-associated suppression of the microbiome–S-equol axis. Cd exposure was also associated with reduced mRNA expression estrogen receptor alpha signaling at the transcriptional level: ERα mRNA was significantly decreased, whereas ERβ mRNA exhibited a similar downward trend without reaching statistical significance (Fig. 2C). Together, these findings suggest that Cd exposure disrupts an S-equol-producing gut microbiome – serum S-equol – hippocampal estrogen receptor axis, linking Cd-induced microbial alterations to reduced availability of a microbiome-derived estrogenic metabolite and diminished estrogen-related signaling.

**Fig. 1.**
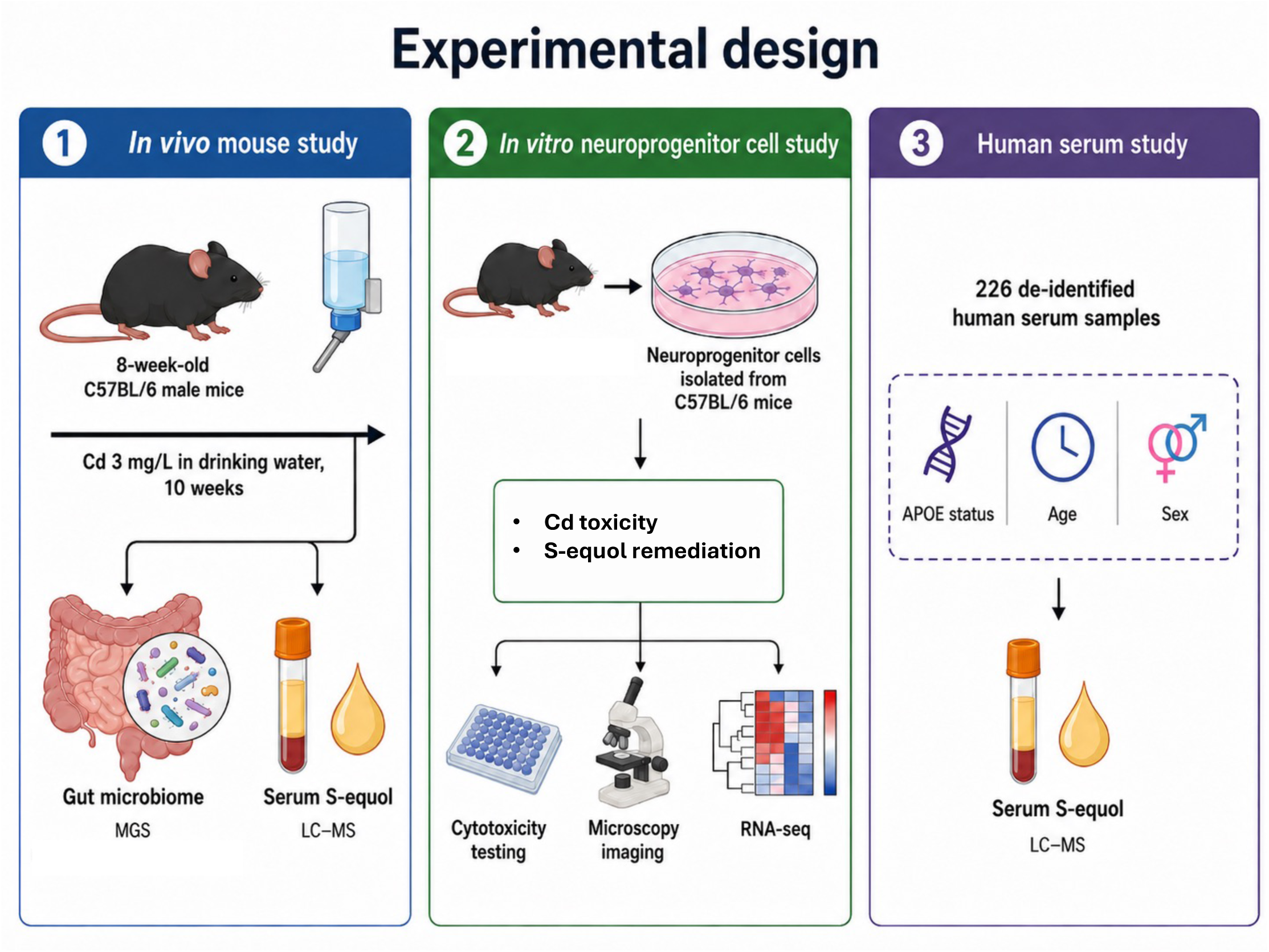
Experimental design integrating *in vivo* mouse study, *in vitro* neural progenitor cell study, and human plasma study. Eight-week-old male C57BL/6 mice were exposed to cadmium (CdCl_2_; 3 mg/L in drinking water) for 10 weeks. Gut microbiome composition was analyzed by metagenomic sequencing (MGS), and circulating S-equol was quantified by liquid chromatography-mass spectrometry (LC-MS). In parallel, neural progenitor cells isolated from C57BL/6 mice were treated with vehicle or Cd with or without S-equol and evaluated by cytotoxicity assays, microscopy imaging, and RNA sequencing (RNA-seq). For translational assessment, plasma S-equol was quantified by LC-MS in 226 de-identified plasma serum samples with available *APOE* status, age, and sex information.

**Fig. 2.**
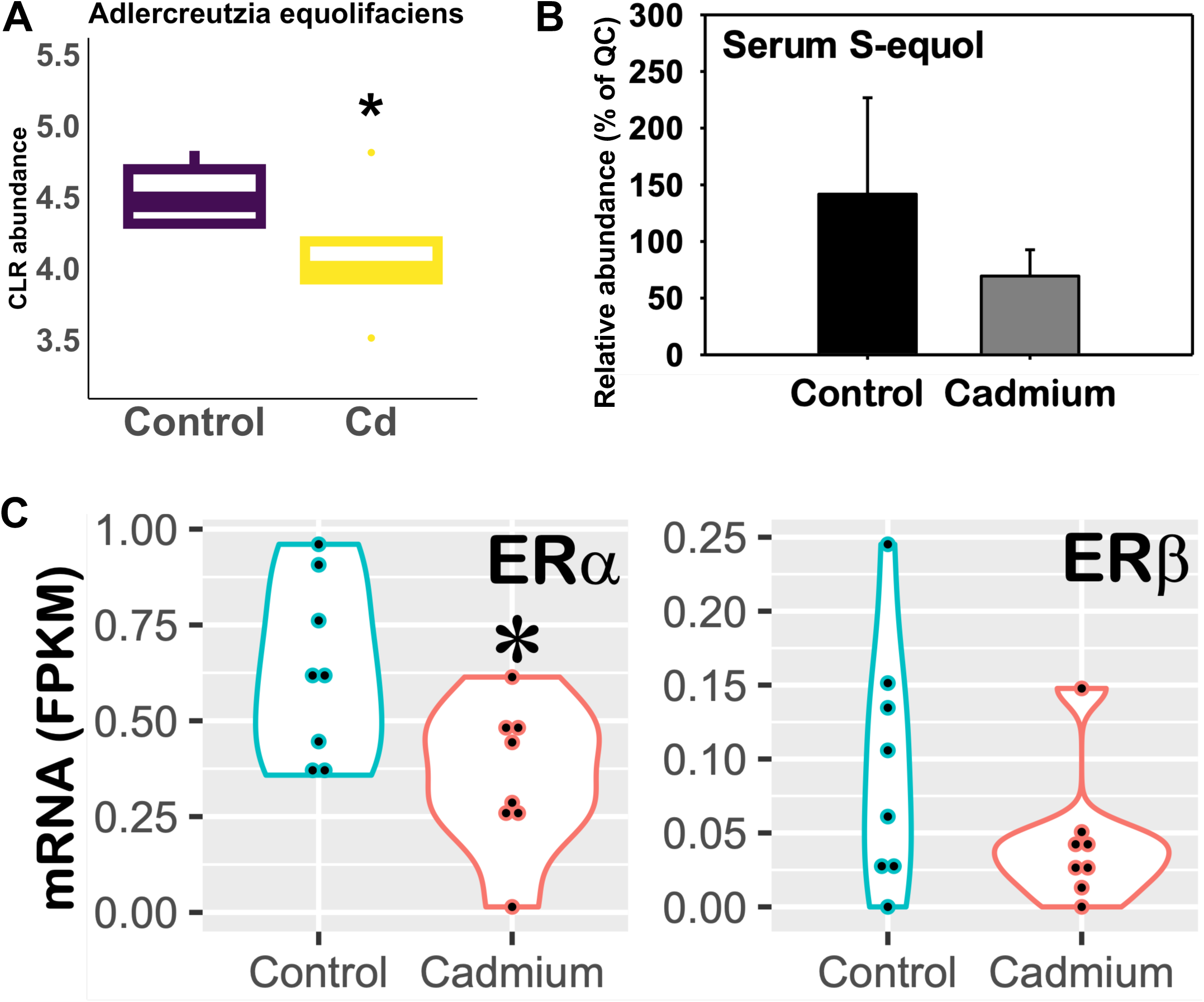
Cadmium exposure disrupts the gut microbiome-S-equol-estrogen receptor axis in mice. **A.** Centered-log ratio normalized (CLR) abundance of the known equol-producing intestinal bacterium *Adlercreutzia equolifaciens* in control and Cd-exposed mice (n=5 per group), showing reduced abundance following Cd exposure. To note, *Adlercreutzia equolifaciens* is also one of the major characterized S-equol-producing bacteria in the human gut^14, 46^. Asterisk (*) represents statistically significant difference as compared to the vehicle group (DESeq2, FDR-BH < 0.05). **B.** Circulating S-equol abundance in control and Cd-exposed mice, demonstrating a decreasing trend following Cd exposure (although a statistical significance was not achieved, Student’s t-test). Plasma S-equol was quantified using LC-MS as described in MATERIALS AND METHODS (n=5 per group). S-equol is reported as relative response, defined as the analyte peak area expressed as a percentage of the pooled quality-control sample, rather than as an absolute concentration. **C.** Messenger RNA expression of estrogen receptor-α (ERα) and estrogen receptor-β (ERβ), showing significantly reduced ERα expression and a non-significant decreasing trend of ERβ expression in Cd-exposed mice (n=8 per group, DESeq2, FDR-BH < 0.05). Together, these findings associate Cd exposure with disruption of an equol-producing gut bacterium, reduced circulating S-equol, and altered estrogen receptor signaling. Expression is diaplyed as FPKM, fragments per kilobase of transcript per million mapped reads.

### S-equol protects neural progenitor cells from Cd cytotoxicity

To determine whether S-equol protects neural progenitor cells against Cd-induced injury, we first characterized the effects of Cd and S-equol individually on cell viability (Fig. 3A-3B). WST-8 viability assay showed that Cd exposure produced a clear concentration-dependent reduction in cell viability, with viability declining to approximately 83% at 5 μM Cd, 74.5% at 7.5 μM, and 68.3% at 10 μM compared with vehicle controls (Fig. 3A). Representative microscopy images were consistent with progressive cellular injury at the higher Cd concentrations. In contrast, S-equol alone showed no appreciable cytotoxicity over the tested range of 10-100 μM, with cell viability remaining near 100% of control (Fig. 3B).

**Fig. 3.**
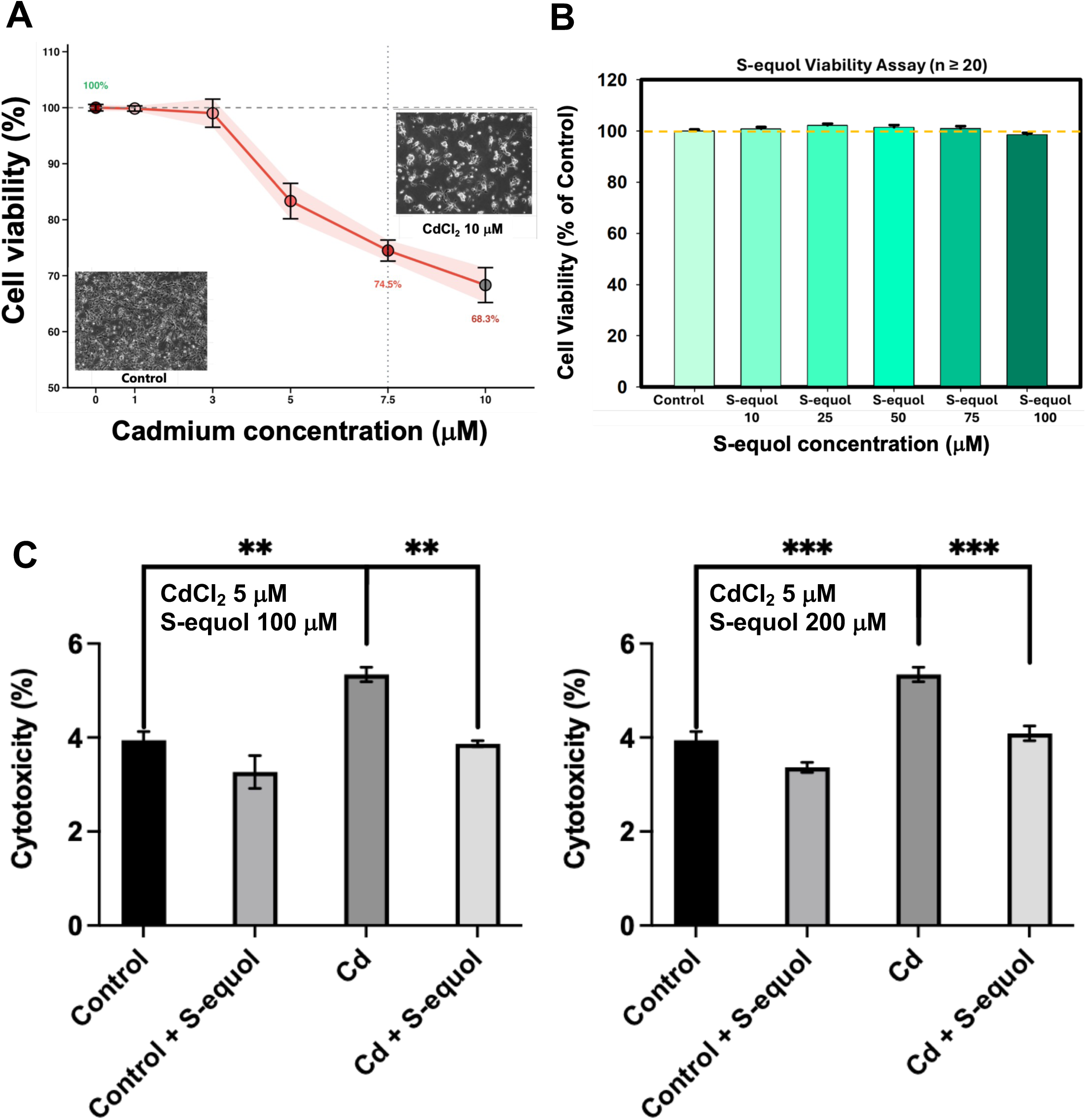
S-equol (100 and 200 μM) attenuates 5 μM Cd-induced cytotoxicity in mouse neural progenitor cells. **A.** Cd dose-response study showing progressive loss of neural progenitor cell viability within increasing Cd concentrations (0, 1, 3, 5, 7.5, 10 μM), with representative microscopy images of control and Cd-treated cells. Cell viability was assessed by quantifying metabolic activity of viable cells, and data were analyzed by one-way ANOVA followed by Dunnett’s test against vehicle control (n=25 control, n=9 per Cd concentration, *p* < 0.05). **B.** S-equol dose-response study showing no apparent toxicity of S-equol on neural progenitor cell viability (one-way ANOVA, *p* < 0.05). **C.** Cytotoxicity in cells treated with Cd (5 μM) with or without S-equol at 100 μM (left) or 200 μM (right). S-equol co-treatment reduced Cd-associated cytotoxicity toward control levels, supporting a protective effect of S-equol against Cd-induced cellular injury (n=5 per group). CytoTOX96 LDH assay was used to quantify LHD released into the culture medium when the plasma membrane is damaged (indicator of cell death). Twenty-four hours after treatment, the CytoTo96 Non-Radioactive Cytotoxicity Assay was used to quantify lactate dehydrogenase (LDH) released from cells into the surrounding medium, which is an indicator of cell death. ** *p* < 0.01; *** *p* < 0.001 (two-way analysis of variance [ANOVA] followed by Tukey’s post hoc test).

We next examined whether S-equol could mitigate Cd-induced cell death using a stepwise dosing strategy, first testing relatively high S-equol concentrations to establish proof-of-concept protection (Fig. 3C). We utilized the lactate dehydrogenase (LDH) cytotoxicity assay, which is a complementary assay to cell viability test by directly quantifying membrane damage. We observed that exposure to 5 μM Cd significantly increased the LDH release which is an indicator of cell death, as compared to the vehicle group. Co-treatment with either 100 or 200 μM S-equol significantly reduced Cd-induced LDH release, indicating partial cyto-protection against death (Fig. 3C, *p* < 0.01 at 100 μM S-equol dose and *p* < 0.001 at 200 μM S-equol dose, respectively). Importantly, S-equol alone at either dose did not increase cytotoxicity. Together, these findings demonstrate that S-equol is well tolerated at the tested concentrations and significantly attenuates Cd-induced cytotoxicity in mouse neural progenitor cells.

### S-equol protects at lower doses under greater Cd stress

Having established proof-of-concept protection at relatively high S-equol concentrations, we next examined whether this effect could be maintained at substantially lower S-equol doses under a greater Cd challenge. Cd was therefore increased from 5 to 7.5 μM, whereas S-equol was reduced to 10 and 25 μM. Despite the greater Cd-induced loss of viability, S-equol remained protective, with 25 μM S-equol significantly improving viability (Fig. 4A). Consistent with reduced cellular stress, Cd strongly up-regulated its prototypical target – the metal-responsive gene metallothionein 2 (*Mt2*), whereas S-equol co-treatment attenuated this response (Fig. 4B). Importantly, the 25 μM represents a substantially lower concentration that nonetheless retained measurable protection against the more stringent 7.5 μM Cd challenge. Together, these findings show that S-equol-mediated protection is maintained at a lower dose under greater Cd toxicity, further strengthening evidence that the remediation effect is not restricted to high S-equol concentrations.

**Fig. 4.**
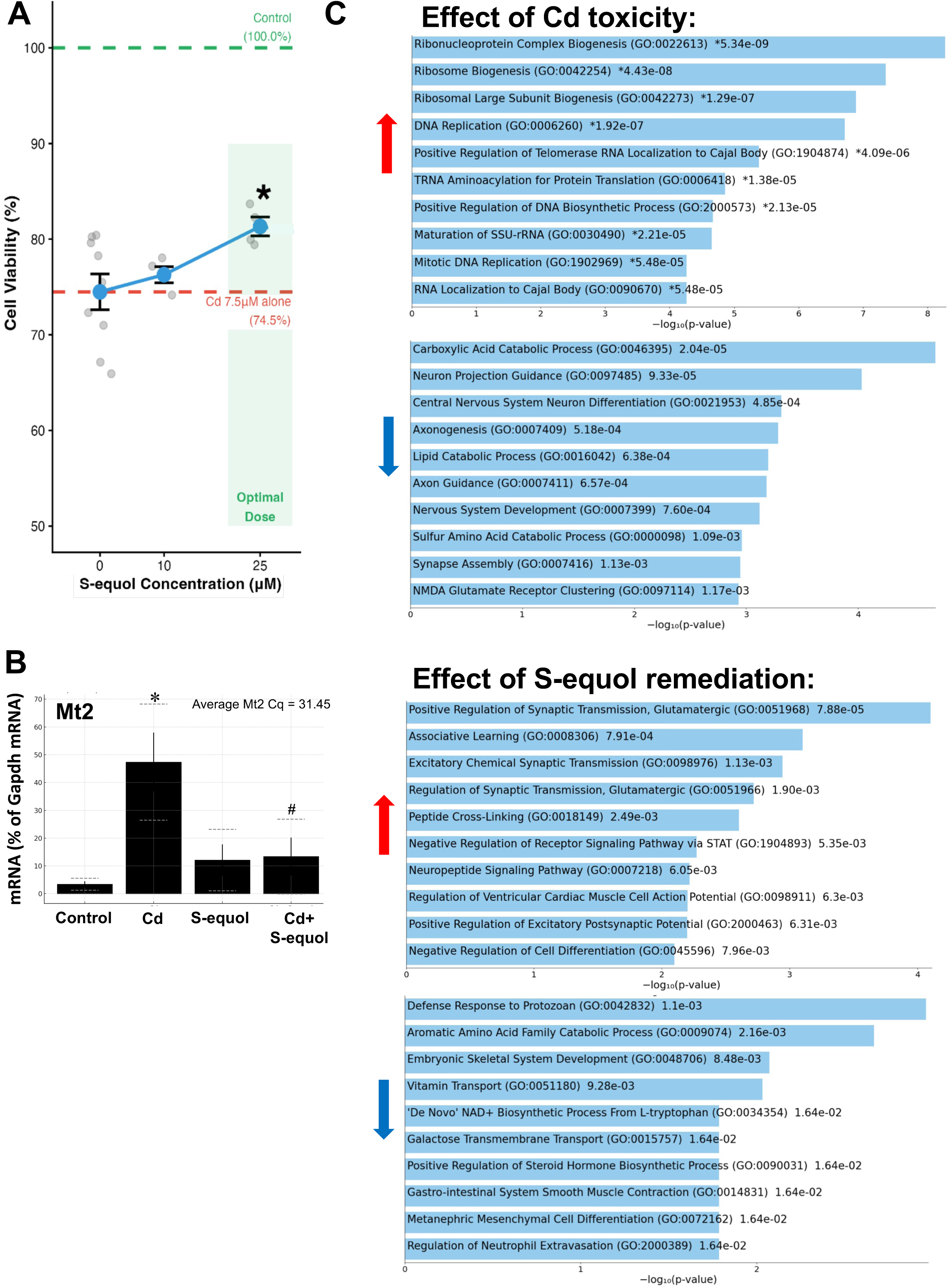
A lower dose of S-equol (25 μM) attenuates cellular stress induced by a higher Cd dose (7.5 μM) and restores neuronal and synaptic transcriptional programs. A. S-equol dose-response under Cd exposure, showing improved neural progenitor cell viability with increasing S-equol concentrations with protection observed at 25 μM S-equol dose against 7.5 μM Cd dose. Asterisk (*) represent statistically significant difference as compared to the Cd alone group (one-way ANOVA followed by Tukey’s *post hoc* test). **B.** Expression of the prototypical Cd-target gene metallothionein 2 (*Mt2*), which is a metal-responsive stress gene, in neural progenitor cells. RT-qPCR was performed in total RNA isolated from the neural progenitor cells (n=5 per group). Asterisk (*) represents statistically significant difference as compared to the vehicle group; pound ((#) represents statistically significant difference as compared to the Cd group. Two-way ANOVA followed by Tukey’s *post hoc* test (*p* < 0.05). **C.** Gene Ontology Biological Process enrichment analysis of RNA-seq data from neural progenitor cells. Cd-associated transcriptional changes included increased ribonucleoprotein and ribosome biogenesis, DNA replication, and biosynthetic programs, together with suppression of neuron projection guidance, central nervous system neuron differentiation, axonogenesis, axon guidance, nervous-system development, synapse assembly, and NMDA glutamate receptor clustering. In contrast, S-equol remediation preferentially enriched neuronal and synaptic processes, including positive regulation of glutamatergic transmission, associative learning, excitatory chemical synaptic transmission, excitatory postsynaptic potential, and neuropeptide signaling. GO analysis was performed using genes meeting *p* < 0.05 and |log₂FC| > 1 (DESeq2).

### S-equol restores neuronal and synaptic gene programs

As a first step to define the molecular programs underlying S-equol-mediated protection, we performed RNA-seq followed by Gene Ontology Biological Process enrichment analysis in neural progenitor cells (Fig. 4C). Cd exposure produced a broad transcriptional shift characterized by increased enrichment of ribonucleoprotein and ribosome biogenesis, DNA replication, and biosynthetic/proliferative processes, together with suppression of pathways central to neuronal development and function, including neuron projection guidance, central nervous system neuron differentiation, axonogenesis, axon guidance, nervous-system development, synapse assembly, and NMDA glutamate receptor clustering (Fig. 4C top two panels).

In contrast, S-equol co-treatment preferentially shifted the transcriptomic response toward restoration of neuronal and synaptic functions. Among the most prominent enriched processes were positive regulation of glutamatergic synaptic transmission, associative learning, excitatory chemical synaptic transmission, regulation of glutamatergic transmission, positive regulation of excitatory postsynaptic potential, and neuropeptide signaling. Conversely, S-equol remediation also downregulated a distinct set of biological processes, including defense response to protozoan, aromatic amino acid family catabolism, vitamin transport, de novo NAD⁺ biosynthesis from L-tryptophan, galactose transmembrane transport, positive regulation of steroid hormone biosynthesis, gastrointestinal smooth muscle contraction, metanephric mesenchymal cell differentiation, and regulation of neutrophil extravasation (Fig. 4C, bottom two panels). These changes suggest that S-equol not only restores neuronal and synaptic transcriptional programs but also suppresses broader metabolic, immune/defense, and non-neuronal differentiation programs associated with the Cd-exposed transcriptional state. Together, the bidirectional pathway changes indicate that S-equol shifts the Cd-disrupted transcriptome toward neuronal and synaptic function while simultaneously reducing metabolic, inflammatory/defense, and non-neuronal programs, consistent with broad transcriptional remediation of Cd-induced cellular stress.

### Plasma S-equol varies with age, APOE4 status and BMI

To explore the translational relevance of S-equol, we examined plasma S-equol in 226 participants from the Arivale cohort, which is a deeply phenotyped longitudinal scientific wellness cohort previously described by Wilmanski *et al.*^49^ in relation to age, apolipoprotein E4 (*APOE*4) status, and body mass index (BMI) (Fig. 5). We quantified cadmium concentrations in these plasma samples using inductively coupled plasma mass spectrometry (ICP-MS), but signals were generally below the analytical detection limit. Because cadmium partitions predominantly into erythrocytes and circulating plasma concentrations are correspondingly low, plasma is not an informative matrix for assessing cadmium body burden; whole blood or urine would be required. Accordingly, associations between circulating Cd and S-equol could not be evaluated in this human dataset.

**Fig. 5.**
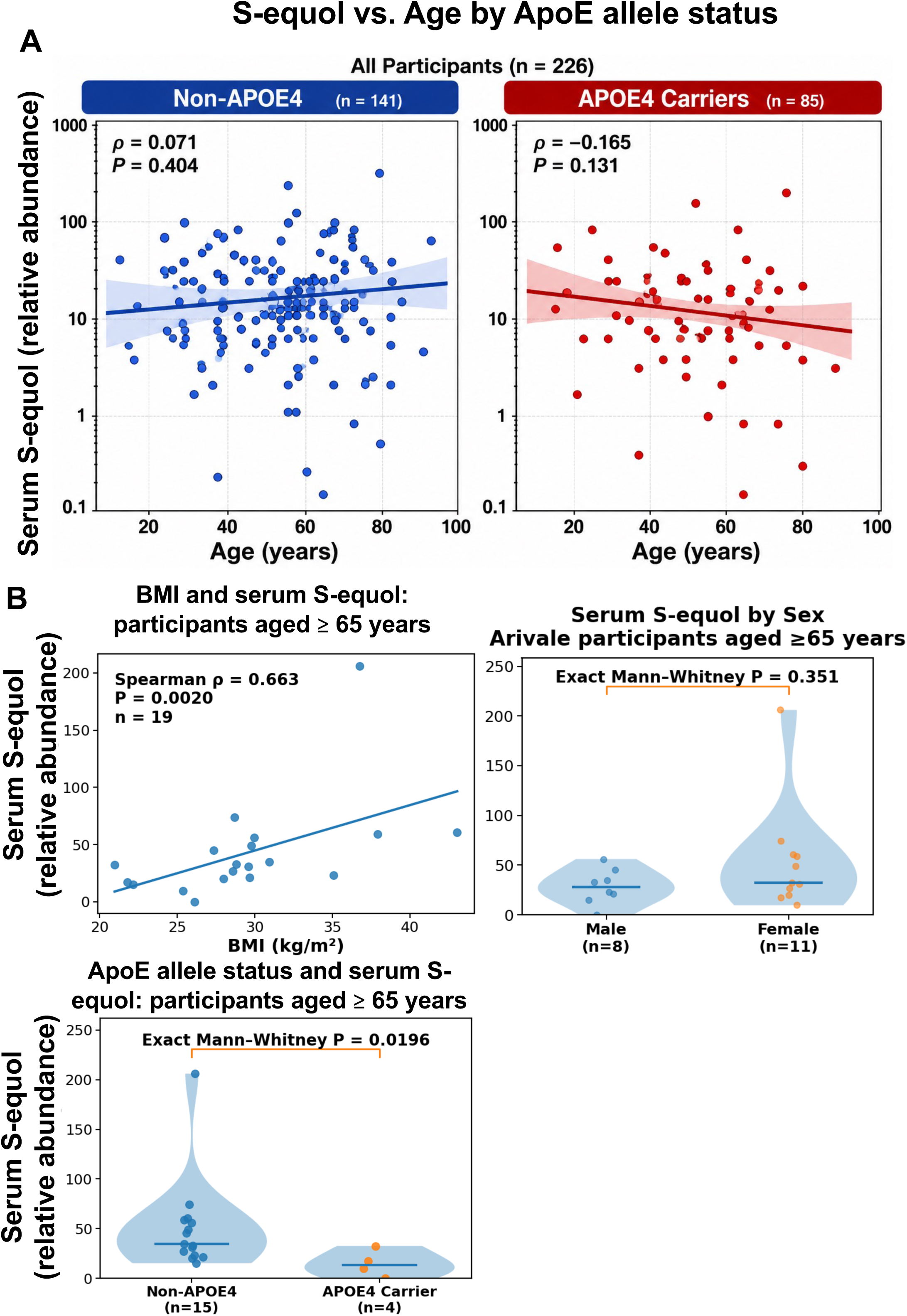
Circulating S-equol in relation to age, *APOE*4 status, and BMI in humans. **A.** Association between age and plasma S-equol among all 226 participants, stratified by *APOE*4 carrier status. No significant age-associated correlation was observed in non-*APOE*4 individuals (n = 141; Spearman ρ = 0.071, *p* = 0.404) or *APOE*4 carriers (n = 85; ρ = −0.165, *p* = 0.131). Lines indicate fitted trends and shaded regions indicate 95% confidence intervals. S-equol is reported as relative response, defined as the analyte peak area expressed as a percentage of the pooled quality-control sample, rather than as an absolute concentration. **B.** Exploratory analyses restricted to participants aged ≥65 years. Plasma S-equol was positively associated with BMI (n = 19; Spearman ρ = 0.663, *p* = 0.0020) and was significantly lower in *APOE*4 carriers (n = 4) than in non-*APOE*4 participants (n = 15; exact Mann– Whitney *p* = 0.0196). S-equol is reported as relative response, defined as the analyte peak area expressed as a percentage of the pooled quality-control sample, rather than as an absolute concentration.

Participant-level metadata, including age, sex, BMI, *APOE* status, and plasma S-equol abundance, are provided in Supplementary Table 1. *APOE*4 is the strongest known genetic risk factor for sporadic/late-onset Alzheimer’s disease, whereas *APOE*3 is the most common allele and is generally used as the reference genotype^10, 50^. Across the full cohort, plasma S-equol was not significantly associated with age in either non-*APOE*4 participants (n = 141; Spearman ρ = 0.071, *p* = 0.404) or *APOE*4 carriers (n = 85; ρ = −0.165, *p* = 0.131); however, the directions of association trend were opposite: S-equol showed a weak tendency to increase with age in non-*APOE*4 participants, whereas it tended to decrease with age in *APOE*4 carriers (Fig. 5A). Because advancing age is the strongest risk factor for AD and late-onset AD is conventionally defined by onset at ≥65 years, we next focused on participants aged ≥65 years to determine whether S-equol associations became more apparent in this higher-risk stage of life. Within this older subgroup, plasma S-equol was positively associated with BMI (Spearman ρ = 0.663, *p* = 0.0020; Fig. 5B). BMI is of interest here for three reasons: adipose tissue acts as a reservoir for isoflavones and their metabolites, with equol detectable in human adipose and glandular tissue; body composition covaries with gut microbial community structure, the upstream determinant of equol production; and adiposity modifies hepatic *Ugt* expression and glucuronidation capacity, the same host-clearance step examined below. Females showed a non-significant trend toward higher plasma S-equol than males (median 32.49 vs. 27.93; exact Mann–Whitney *p* = 0.351). Although not statistically significant, this directional difference suggests that sex may contribute modestly to S-equol variability in older adults. In addition, *APOE*4 carriers exhibited significantly lower plasma S-equol than non-*APOE*4 participants (exact Mann-Whitney *p* = 0.0196). These findings suggest that relationships between circulating S-equol, metabolic status, sex, and *APOE*4 genotype may become more apparent in older adults. Given the small number of individuals aged ≥ 65 years, however, these subgroups observations should be considered exploratory and warrant validation in larger cohorts.

### APOE genotype and sex alter hepatic Ugt expression in mice

Because circulating S-equol concentrations reflect not only microbial production but also host metabolism and clearance, we examined hepatic expression of UDP-glucuronosyltransferase (UGT) genes in humanized *APOE*3 and *APOE*4 transgenic mice at 75-weeks of age, which is equivalent to humans in their early 60s. To note, S-equol is generated by gut microbial conversion of the soy isoflavone daidzein and, after absorption, undergoes extensive phase II metabolism, predominantly glucuronidation in the liver^24, 27, 51^. Glucuronidation reduces the circulating free S-equol pool and facilitates urinary or biliary elimination; glucuronide conjugates constitute the predominant circulating and excreted forms of S-equol^27, 52^. Thus, we hypothesize that *APOE*4-associated induction of S-equol-metabolizing UGT pathways could increase S-equol glucuronidation and clearance, thereby contributing to the lower circulating S-equol observed in older human *APOE*4 carriers.

In livers of aged humanized *APOE*3 and *APOE*4 transgenic mice, the hepatic expression profiles shown here reveal marked sex-dependent differences in multiple *Ugt* genes (Fig. 6), with substantially lower expression in females for several *Ugt*s. In addition, *APOE* genotype-dependent effects are evident for selected *Ugt* genes, particularly in males, where *APOE*4 mice showed higher hepatic expression than *APOE*3 mice. These findings raise the possibility that *APOE* genotype and sex influence circulating S-equol not only through upstream microbial production but also through downstream hepatic conjugation and clearance.

**Fig. 6.**
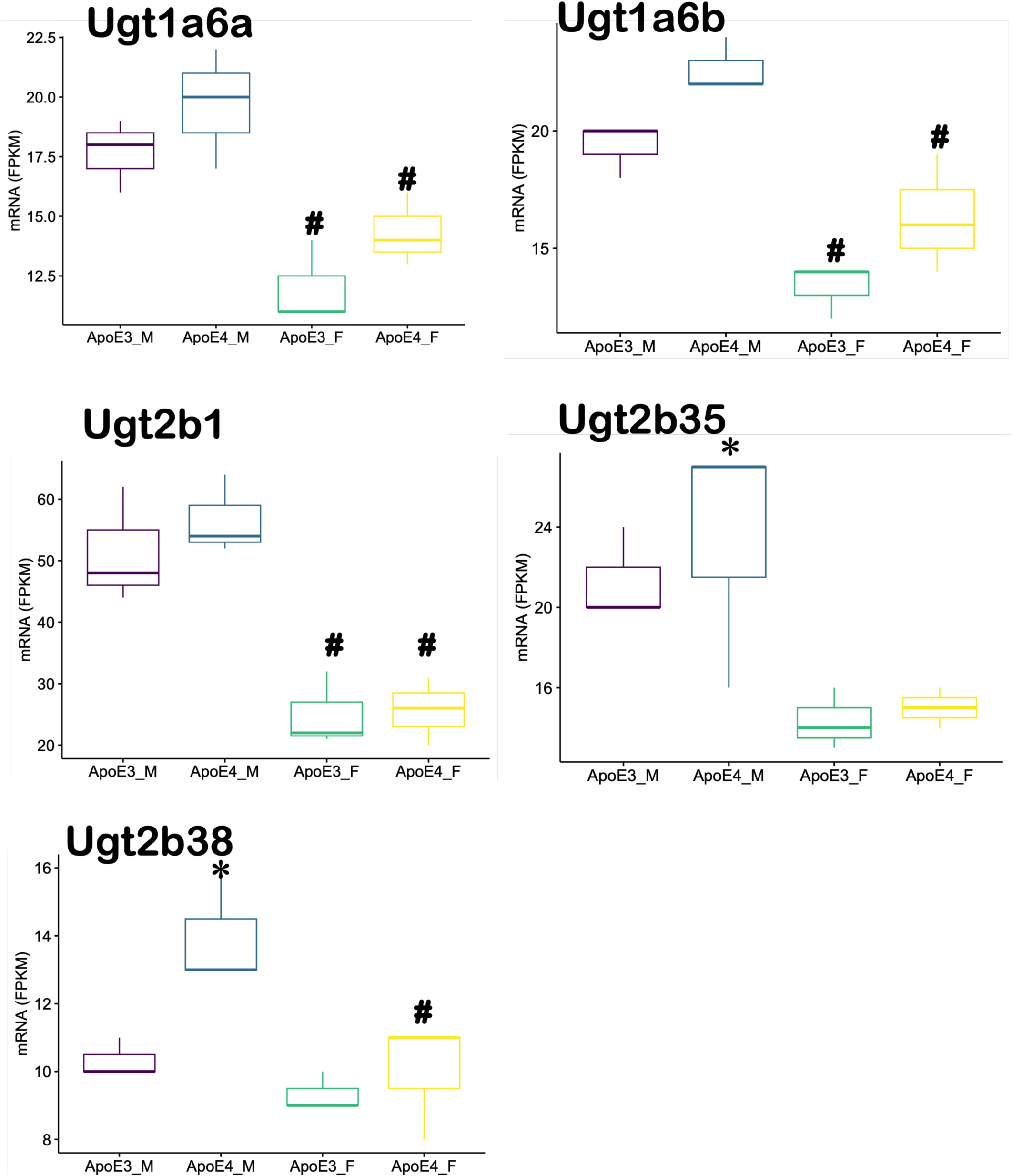
Hepatic expression of differentially regulated *Ugt* genes in aged humanized *APOE*3 and *APOE*4 mice. Messenger RNA expression of distinct *Ugt* genes was quantified by RNA-seq in livers of 75-week-old male and female humanized *APOE*3 and *APOE*4 transgenic mice (n = 3 per group). Asterisks (*) indicate a significant *APOE* genotype effect and pounds (#) indicate a significant sex effect (*p* < 0.05; two-way ANOVA followed by Tukey’s honestly significant difference [HSD] test).

## DISCUSSION

Our findings identify disruption of a gut microbiome-S-equol-brain-estrogen signaling axis as a potential mechanism linking Cd exposure to neurological vulnerability. Cd is known to alter gut microbial composition and metabolic function^53, 54^, and our previous studies showed that Cd disrupts the gut-liver axis in humanized *APOE*3/E4 mice^9^ and that gut dysbiosis precedes hippocampus-dependent learning and memory deficits^12^. Here, Cd significantly reduced *Adlercreutzia equolifaciens*, a well-characterized human intestinal equol-producing bacterium, accompanied by a downward trend in circulating S-equol. This is biologically relevant because S-equol preferentially activates ERβ and has been linked experimentally to enhanced mitochondrial bioenergetics and neuroprotective, synapse-preserving effects. The concurrent reduction in hippocampal ERα expression and downward trend in ERβ therefore raise the possibility that Cd perturbs both the microbial supply of an estrogenic metabolite and its neural signaling pathway. However, these findings remain associative: circulating S-equol did not decrease significantly, receptor expression does not establish receptor activity, and *A. equolifaciens* abundance alone does not prove equol-producing capacity because strain-level variation exists^55^. Thus, microbiome manipulation and *in vivo* S-equol replacement studies in the future will be required to establish causality.

A second major finding is that S-equol directly mitigates Cd-induced neural injury and shifts the transcriptional response toward restoration of neuronal and synaptic function. Cd produced a dose-dependent loss of neural progenitor-cell viability, increased LDH release, and strongly induced Mt), a prototypical metal-responsive stress gene that is expressed in the brain, including the hippocampus, and is inducible by Cd and other metals^56, 57, 58, 59^. In contrast, S-equol reduced cytotoxicity, improved viability, and attenuated *Mt2* induction; notably, protection remained detectable at 25 μM S-equol under a higher Cd challenge. These findings are consistent with previous evidence that Cd impairs neural progenitor-cell survival, proliferation, and differentiation through stress-responsive signaling pathways^8, 60^. Transcriptomic analysis further showed that Cd suppressed programs involving neuronal differentiation, axonogenesis, synapse assembly, and NMDA receptor organization, whereas S-equol preferentially restored glutamatergic transmission, excitatory postsynaptic signaling, neuropeptide signaling, and learning-associated pathways. This pattern is biologically plausible given prior evidence that S-equol or equol protects neurons from neurotoxic insults^20^, prevents synaptic loss through an ERβ-dependent mechanism^23^, protects against Aβ-induced neuronal injury through estrogen-receptor signaling^22^, promotes neurite and dendritic development^61^, and enhances mitochondrial respiration and bioenergetic capacity in experimental neural systems^21^. Collectively, these data suggest that S-equol acts beyond simple preservation of cell viability by shifting the Cd-disrupted cellular state toward neuronal and synaptic function. However, the transcriptomic findings identify pathways associated with protection rather than causal mechanisms, and direct measurements of mitochondrial function, oxidative stress, synaptic activity, and ERβ dependence will be required in future studies to define the mechanisms responsible for S-equol-mediated protection. A further limitation is that the S-equol concentrations used here (10 – 200 μM) substantially exceed those achievable in humans. In equol producers, circulating total S-equol following a soy challenge is typically 10-140 nmol/L, producer status is conventionally defined at concentrations above 83 nmol/L, and less than 1% circulates as the unconjugated aglycone. These experiments therefore establish biological plausibility and identify candidate mechanisms rather than demonstrating physiological sufficiency, and lower-concentration or repeated-exposure paradigms will be required to assess translational relevance.

Another major finding is that S-equol bioavailability may be shaped not only by microbial production but also by sex- and *APOE*-dependent host metabolism. Following absorption, S-equol undergoes extensive phase-II conjugation, predominantly glucuronidation, with hepatic microsomes displaying greater S-equol glucuronidation capacity than intestinal microsomes^24^. In humans, <1% of circulating S-equol after oral administration is present in the unconjugated form^25^, and approximately 82% of an administered dose can be recovered in urine, consistent with efficient conjugation and elimination^26^; glucuronide conjugates represent major circulating and excreted S-equol metabolites^27, 52^. Thus, hepatic UGT activity represents an important downstream determinant of exposure to biologically available S-equol. Circulating S-equol may also be shaped by microbial deconjugation. Although renal excretion of S-equol glucuronide is the dominant elimination route, biliary secretion delivers conjugated isoflavones to the colon, where microbial β-glucuronidases can regenerate the free aglycone for reabsorption, and pharmacokinetic studies of equol have reported secondary plasma peaks consistent with enterohepatic recycling. Recent characterization of distinct gut microbial β-glucuronidase subtypes capable of reactiving conjugated endobiotics therefore raises the possibility that the gut microbiome influences S-equol exposure at both the production and recycling steps. Consistent with established sex-dependent regulation of hepatic *Ugt* genes in mice^62, 63^ and sex differences in selected *UGT* expression and glucuronidation activity in human liver^64, 65^, we observed pronounced sex-dependent expression of multiple hepatic *Ugt* genes in aged humanized *APOE*3/E4 mice, together with genotype effects in selected *Ugt*s, particularly higher expression in *APOE*4 than *APOE*3 males. These findings raise the possibility that the lower circulating S-equol observed in older human *APOE*4 carriers may reflect not only differences in microbial production but also enhanced downstream conjugation and clearance. Future studies integrating S-equol pharmacokinetics with UGT activity and metabolite profiling across sex and *APOE* genotype will provide additional insights into this proposed host-control mechanism.

The present study also took an initial step in characterizing the age- and *APOE* allele-modified pattern of circulating S-equol in humans. *APOE*4 is the strongest established genetic risk factor for sporadic AD, and among participants aged ≥65 years in our cohort, *APOE*4 carriers exhibited significantly lower plasma S-equol than non-carriers, while S-equol was positively associated with BMI; females also showed a non-significant tendency toward higher S-equol than males. These observations are notable given the inconsistent literature relating soy intake itself to cognitive outcomes: long-term soy-isoflavone supplementation did not improve global cognition in postmenopausal women^34^, and a large Japanese cohort found no association between total soy or isoflavone intake and disabling dementia overall^33^. In contrast, studies focusing specifically on equol exposure or equol-producing status provide evidence of potential neurological relevance. In patients with AD, soy isoflavone supplementation produced no overall cognitive benefit, but higher plasma equol was associated with better verbal fluency and speeded dexterity in exploratory analyses^40^; among cognitively normal older Japanese adults, equol-producing status was inversely associated with cerebral white-matter lesion burden^41^; and a small S-equol intervention study in women with AD demonstrated preliminary mitochondrial target engagement^42^. These findings support the possibility that circulating S-equol, rather than soy exposure alone, may better capture the biologically relevant interaction between diet and the gut microbiome. Indeed, equol-producing capacity varies markedly among individuals and is influenced by microbial and dietary factors^37, 38, 66^. Importantly, our human observations remain hypothesis-generating: the ≥65-year subgroup was small (n = 19), including only four *APOE*4 carriers, the sex difference was not statistically significant, and Cd concentrations were largely below detection, which does not allow direct evaluation of a Cd–S-equol relationship in this cohort. Thus, larger prospective cohorts with simultaneous measurements of Cd exposure, equol-producing status, circulating free and conjugated S-equol, *APOE* genotype, and cognitive outcomes will be required to establish the clinical relevance of this interaction. The ongoing 369-participant randomized ACE trial directly testing S-equol effects on vascular and cognitive aging sill further underscore the translational importance of this question^43^.

Collectively, our findings support a model, summarized in Fig. 7, in which environmental exposure, gut microbial metabolism, host genetics, sex, and hepatic biotransformation converge to shape the availability of a neuroactive microbial metabolite and, potentially, individual susceptibility to neurotoxicity. This framework extends our previous observations that Cd disrupts the gut-liver axis in an *APOE*-dependent manner^9^ and that Cd-induced gut dysbiosis precedes hippocampus-dependent cognitive deficits^12^. In the present study, Cd disrupted the *A. equolifaciens*-S-equol-estrogen receptor axis, whereas S-equol directly attenuated Cd-induced cytotoxicity and shifted neural progenitor-cell transcription toward neuronal, glutamatergic, synaptic, and learning-related functions. These findings are particularly relevant to aging and *APOE*4, the strongest genetic risk factor for sporadic AD^10^, because equol-producing status has been associated with lower cerebral white-matter lesion burden in older adults^41^, and S-equol has shown ERβ-dependent synaptic protection and preliminary mitochondrial target engagement in experimental and human AD studies^23, 42^. Nevertheless, several important questions remain: the present study does not establish whether loss of *A. equolifaciens* causes reduced S-equol availability, whether altered hepatic *Ugt* expression changes S-equol pharmacokinetics, or whether S-equol supplementation prevents Cd-induced cognitive impairment *in vivo*. Moreover, the Cd-exposed mouse study was restricted to males because we previously showed that male mice were more susceptible to Cd neurotoxicity^67^, and the human ≥65-year subgroup was small and lacked quantifiable Cd exposure. Future studies incorporating microbiome manipulation, measurement of free and conjugated S-equol, sex- and *APOE*-stratified pharmacokinetics, ERβ perturbation, and behavioral endpoints will be required to establish causality. These studies will be addressed in future directions. Despite these limitations, our findings have provided important insights that microbiome-derived metabolites such as S-equol may represent modifiable determinants of host resilience to environmental neurotoxicants, providing a conceptual framework for precision interventions at the intersection of the exposome, microbiome, and genetic susceptibility.

**Fig. 7.**
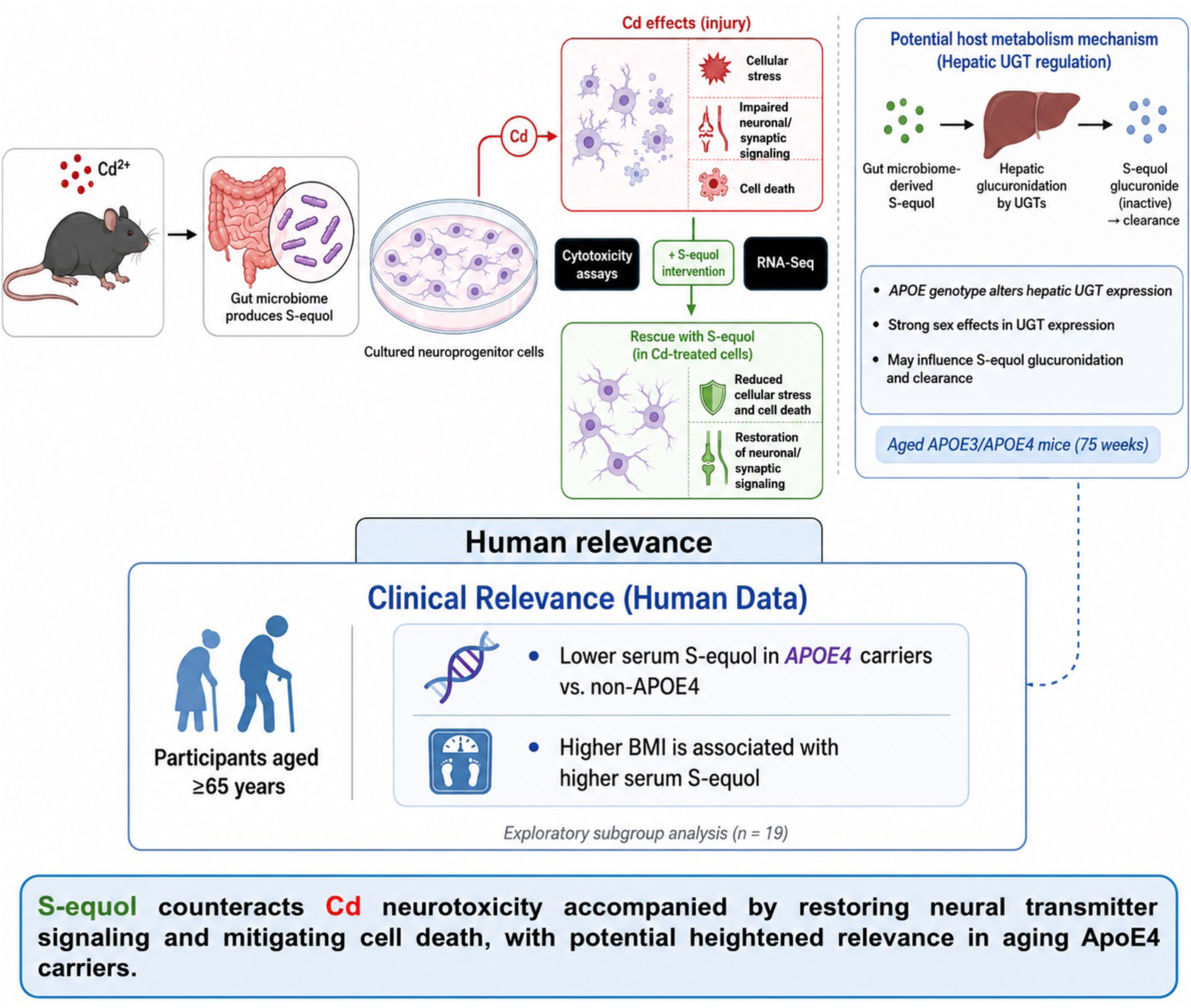
Proposed model linking Cd exposure, microbiome-derived S-equol, and neurotoxicity. Graphical summary of the proposed mechanism and remediation pathway. Cd exposure is associated with decreased abundance of the equol-producing bacterium *Adlercreutzia equolifaciens*, reduced circulating S-equol, and diminished estrogen receptor signaling, accompanied by cellular stress, impaired neuronal and synaptic programs, and neurotoxicity. S-equol intervention attenuates Cd-induced cytotoxicity and cellular stress while restoring neuronal, glutamatergic, synaptic, and learning-related transcriptional programs, resulting in partial remediation of Cd neurotoxicity. Human analyses provide exploratory translational support, suggesting lower circulating S-equol in older *APOE*4 carriers and a positive relationship between S-equol and BMI among individuals aged ≥65 years.

## Methods

### Chemicals and reagents

Cadmium chloride (CdCl_2_, anhydrous; catalog number: 202980; ≥ 99.99% trace metals basis; CAS 10108-64-2; formula weight 183.32) was purchased from MilliporeSigma (Burlington, MA). S-equol was purchased from Sigma Alderich (catalog no. SML2147; ≥ 97% by HPLC; CAS 531-95-3; formula weight 242.27). Dimethyl sulfoxide (DMSO) was used as the vehicle for both compounds and prepared fresh from powder at the time of use. CdCl_2_ powder and pure DMSO were stored at room temperature and S-equol powders and S-equol stock solutions in DMSO were stored at -20 ^0^C. Working solutions were prepared by dilution into culture medium immediately before each exposure. The final DMSO concentration was 0.1% (v/v) and was matched across all treated and vehicle-control wells. Unless otherwise noted, all other chemicals were purchased from Sigma-Aldrich.

### Animal studies

Six-week-old male C57BL/6J mice were obtained from Jackson Laboratories (Bar Harbor, ME) and housed in groups of up to five animals per cage under a 12-h light/12-h dark cycle. Food (PicoLab Rodent Diet 20, LabDiet, St. Louis, MO) and drinking water were provided *ad libitum*. Drinking water consisted of reverse-osmosis-purified tap water that was acidified with 2.4-2.8% HCl and autoclaved. Starting at 8-weeks of age, mice were assigned to receive either control drinking water or water containing 3 mg/L cadmium as CdCl₂ for 9 weeks (**Fig. 1**). At the terminal time point, large intestinal content and hippocampus were collected, frozen in liquid nitrogen, and then stored in -80 °C until further analysis. All procedures involving hazardous materials were conducted in accordance with University of Washington Environmental Health and Safety guidelines, and animal experiments were performed under Institutional Animal Care and Use Committee approval of University of Washington (4339-01). In a separate experiment, RNA-Seq (n=3 per sex per genotype) was conducted in 75-week-old aged humanized *APOE*3 and *APOE*4 knock-in (*APOE*3-KI and *APOE*4-KI) mice in C57BL/6 background, as we described before^9, 68^.

### Fecal DNA extraction and shotgun metagenomic sequencing

Genomic DNA was isolated from the large intestinal content of mice using the AllPrep® PowerFecal DNA/RNA Kit (Qiagen, Hilden, Germany) following the manufacturer’s instructions. DNA yield was measured with a Qubit 2.0 Fluorometer (Life Technologies, Grand Island, NY), and DNA quality and integrity were assessed using an Agilent 2100 Bioanalyzer (Agilent Technologies, Santa Clara, CA). Shallow shotgun metagenomic sequencing was performed by Diversigen (New Brighton, MN).

Sequence reads were mapped against the Diversigen Venti reference database, which includes representative bacterial genomes from RefSeq together with metagenome-assembled genomes (MAGs) and cultured isolate genomes. Only high-quality MAGs with >90% completeness and <5% contamination, as determined by CheckM, were included. Reads were aligned to reference genomes at 97% sequence identity using the fully gapped BURST aligner. When multiple references produced equivalent best matches, assignments were resolved by minimizing the total number of unique operational taxonomic units (OTUs). Taxonomic classification was assigned to the lowest common ancestor supported by at least 80% of equally scoring reference sequences and was based on Genome Taxonomy Database release 95 (GTDB r95). Samples containing fewer than 10,000 sequence reads were excluded. Low-abundance OTUs representing less than one-millionth of all strain-level markers were removed, as were species with <0.01% coverage of unique genomic regions and <0.1% coverage of the complete genome. To account for the compositional nature of microbiome count data, taxonomic abundance profiles were transformed using centered log-ratio (CLR) transformation prior to downstream statistical analyses.

### LC-MS of S-equol quantification

S-equol was quantified by liquid chromatography coupled with tandem mass spectrometry (LC-MS/MS) in serum from the mouse study and in plasma from the human cohort. For sample preparation, frozen samples were thawed overnight at 4 °C. An aliquot of 50 μL from each sample was transferred into a 2 mL Eppendorf tube for protein precipitation and metabolite extraction. Each aliquot was mixed with 500 μL of methanol (MeOH) and 50 μL of an internal standard solution containing 1,810.5 μM ^13^C_3_-lactate and 142 μM ^13^C_5_-glutamic acid. Samples were vortex-mixed for 10 s and incubated at −20 °C for 30 min. Following incubation, samples were centrifuged at 14,000 rpm for 10 min at 4 °C. A 450 μL aliquot of the resulting supernatant was transferred to a clean Eppendorf tube and evaporated to dryness using a CentriVap Concentrator (Labconco, Fort Scott, KS). The dried extracts were subsequently reconstituted in 150 μL of 40% phosphate-buffered saline (PBS)/60% acetonitrile (ACN). A pooled quality-control (QC) sample was prepared by combining aliquots from all study samples. Targeted profiling was performed using a globally optimized targeted mass spectrometry workflow as previously described.

Targeted metabolomic analysis was conducted using an Agilent 1290 UPLC system coupled to an Agilent 6495 triple-quadrupole mass spectrometer (QQQ-MS; Agilent Technologies, Santa Clara, CA). Chromatographic separation was achieved by hydrophilic interaction liquid chromatography (HILIC) using a Waters XBridge BEH Amide column (150 × 2.1 mm, 2.5 μm particle size; Waters Corporation, Milford, MA). The mobile-phase flow rate was maintained at 0.3 mL/min. The autosampler and column compartment temperatures were maintained at 4 °C and 40 °C, respectively. Mobile phase A consisted of 10 mM ammonium acetate and 10 mM ammonium hydroxide in 95% H_2_O/5% ACN, whereas mobile phase B contained 10 mM ammonium acetate and 10 mM ammonium hydroxide in 95% ACN/5% H_2_O. The chromatographic gradient began with an isocratic hold at 90% B for 1 min, followed by a gradual decrease in B to 40% at 11 min. The mobile phase was maintained at 40% B for an additional 4 min, until 15 min, after which B was returned to 90% to re-equilibrate the column before the subsequent injection. The mass spectrometer was operated with an electrospray ionization (ESI) source, and targeted compounds were monitored using multiple-reaction-monitoring (MRM) acquisition. Instrument operation and LC-MS/MS data acquisition were controlled using Agilent MassHunter Workstation software. Extracted MRM chromatographic peaks were integrated using Agilent MassHunter Quantitative Data Analysis software (Agilent Technologies, Santa Clara, CA).

### RT-qPCR

Real-time quantitative polymerase chain reaction (RT-qPCR) was performed using total RNA isolated from neural progenitor cells. Total RNA was reverse-transcribed into single-stranded cDNA using the High-Capacity cDNA Reverse Transcription Kit (Applied Biosystems, Foster City, CA, USA; 1001073) according to the manufacturer’s instructions. The resulting cDNA was amplified using SsoAdvanced Universal SYBR Green Supermix on a Bio-Rad CFX384 Real-Time PCR Detection System (Bio-Rad, Hercules, CA, USA). Primers were synthesized by Integrated DNA Technologies (IDT; Coralville, IA, USA). Relative gene expression was determined using the ΔΔCq method. Expression of mouse metallothionein 2 (*Mt2*) was normalized to the housekeeping gene glyceraldehyde-3-phosphate dehydrogenase (Gapdh) and expressed relative to the corresponding control group. The mouse *Mt2* primer sequences are: forward, 5′-TGTGCTGGCCATATCCCTTG-3′; reverse, 5′-GCGGAGAGTATTGGGTCGAG-3′. The mouse Gapdh primer sequences are: forward: 5’-GGCAAATTCAACGGCACAGT-3’; reverse: 5’-GTCTCGCTCCTGGAAGATGG-3’.

### Isolation and treatment of neuroprogenitor cells

Primary adult neural progenitor cells (aNPCs) were isolated from the subgranular zone (SGZ) of the dentate gyrus of 6.5-9-week-old male C57BL/6 mice as previously described^8^. SGZ tissue was micro-dissected, enzymatically and mechanically dissociated, filtered, and enriched by Percoll centrifugation. Cells were cultured in Neurobasal-based proliferation medium supplemented with B27, L-glutamine, penicillin/streptomycin, EGF, and bFGF at 37°C and 6.5% CO₂. Primary neurospheres formed within 7–14 days, were dissociated, and maintained in growth medium containing N2, B27, heparin, EGF, and bFGF. Cultures were passaged no more than 11 times.

### Cell viability assay

Cell viability was assessed using the Cell Viability Assay kit (Colorimetric; ab112118; Abcam, Waltham, MA), in which a water-soluble dye shifts its absorption spectrum on reduction by metabolically active cells. After the 24 h exposure, assay solution was added to each well and plates were incubated for 4 hr at 37 ^0^C. Absorbance was read at 570 and 605nm on a SpectraMax i3x reader (molecular Devices, San Jose, CA). Statistical analyses were performed in R v.4.5.2. Cell viability was normalized to the mean of vehicle-treated wells on the same plate and is presented as mean ± s.d. For the cadmium (Cd) dose-response viability assay, differences among treatment groups were analyzed by one-way ANOVA followed by Dunnett’s multiple-comparisons test against the vehicle control (*n* = 25 for vehicle; *n* = 9 per Cd concentration). Because the S-equol dose-response experiment was designed to assess whether S-equol alone produced a biologically meaningful change in viability, equivalence to vehicle control was evaluated using two one-sided tests (TOST) with prespecified equivalence bounds of ±10%, rather than inferring equivalence from a non-significant difference (*n* = 25 for vehicle; *n* = 20 per S-equol concentration). For Cd–S-equol combination experiments, within each Cd concentration, combination treatments (*n* = 4) were compared with the corresponding Cd-only treatment (*n* = 9) by one-way ANOVA followed by Dunnett’s multiple-comparisons test. Because S-equol alone was equivalent to vehicle control within the prespecified bounds, Bliss independence predicts combined viability equivalent to that observed with Cd alone; viability significantly exceeding this expectation was therefore interpreted as evidence of non-additive protection by S-equol.

### Cytotoxicity assay

Twenty-four hours after treatment, cytotoxicity was assessed using the CytoTox 96® Non-Radioactive Cytotoxicity Assay (Promega; catalog no. G1780; Madison, WI) by lactate dehydrogenase (LDH) released from damaged cells into the culture medium as an indicator of cell death. LDH activity was quantified according to the manufacturer’s instructions and compared among treatment groups (n=5 per group. Statistical significance was determined by two-way ANOVA (*p* < 0.01; \**p* < 0.001).

### Total RNA isolation

Total RNA was extracted from hippocampal tissue collected from control and Cd-exposed mice for the *in vivo* study, as well as well as control and treated neural progenitor cells for the *in vitro* study, using RNA-Bee reagent (Tel-Test Inc., Friendswood, TX) following the manufacturer’s instructions. RNA concentration was determined by absorbance at 260 nm using a NanoDrop 1000 Spectrophotometer (Thermo Scientific, Waltham, MA). RNA integrity was assessed by examining 28S and 18S rRNA bands on formaldehyde-agarose gels and was further evaluated with an Agilent 2100 Bioanalyzer by Novogene. RNA sequencing was performed by Novogene (Sacramento, CA) using five biological replicates per group.

### RNA-sequencing and differential expression analysis

For the in vitro neural progenitor cells study, total RNA was isolated and RNA sequencing was performed by Novogene using five biological replicates per group. The cDNA libraries were prepared following poly (A) selection and sequenced on the DNBSEQ platform in paired-end mode (2 x 150 bp), generating approximately 20 million read pairs (6 Gb) per sample. Paired-end reads were aligned to the mouse reference genome GRCm39 (NCBI assembly GCF_000001635.27) using STAR v2.7.11b. Gene-level read counts were quantified with featureCounts^69^ from the Subread package v2.1.1 against the GCF_000001635.27_GRCm39_genomic.gft annotation, in paired-end mode at the exon level summarized by gene_id using 8 threads. Differential gene expression analysis was performed in R v4.5.2 using DESeq2 v1.50.2^70^. Genes meeting the specified significance threshold were used for downstream Gene Ontology enrichment analysis (*p* < 0.05 and |log_2_ fold change| > 1).

### Human plasma samples and statistical analysis

De-identified serum samples from 226 participants in the Arivale cohort were analyzed. The Arivale cohort comprises adults enrolled in a longitudinal scientific wellness program and has been previously described^49^. Available metadata for the present study included age, sex, BMI, and *APOE* status. Plasma S-equol was quantified by LC-MS. Blood was collected by trained phlebotomists during routine clinical draws and centrifuged within 1 h of collection. The plasma layer was separated and held at -20 ^0^C until shipment to the biorepository on dry ice, where samples were thawed once for aliquoting and transferred to -80 ^0^C. All samples were subsequently maintained at -80 ^0^C until analysis and underwent identical collection, processing, and storage procedures. Participants were classified as *APOE*4 carriers or non-carriers. Associations between age and S-equol were evaluated within *APOE*4 strata using Spearman rank correlation. Exploratory analyses were restricted to participants aged ≥65 years; associations between BMI and S-equol were assessed by Spearman correlation, and S-equol abundance was compared between *APOE*4 carriers and non-carriers using an exact two-sided Mann–Whitney U test. Statistical significance was defined as *p* < 0.05. The original Arivale study was conducted under Western Institutional Review Board approval (Study Nos. 20170658).

## Data Availability

All raw and analyzed data will be deposited in the NCBI Gene Expression Omnibus (GEO) database.

## Code Availability

No custom code was generated in this study. All analyses were performed using publicly available software as described in Methods: STAR v2.7.11b, featureCounts from Subread v2.1.1, and DESeq2 v1.50.2 in R v4.5.2

## Author Contributions

Y.S. conducted experiments, performed data analysis, and contributed to manuscript preparation, review, and editing. H.W. conducted experiments, performed data analysis, and contributed to manuscript review and editing. L.L. conducted experiments, performed data analysis, and contributed to manuscript review and editing. H.G. conducted experiments, performed data analysis, and contributed to manuscript review and editing. J.J.-M. performed data analysis and contributed to manuscript review and editing. A.T.M. and S.M.G. provided sample access, contributed to data analysis, and participated in manuscript review and editing. J.Y.C. conceived and designed the study, acquired funding, supervised the work, and led manuscript writing, review, and editing.

## Acknowledgements

The authors would like to thank members of the Cui Laboratory for proof-reading the manuscript.

This work was supported by the National Institute of Environmental Health Sciences (NIEHS) R00ES034068 and P30ES017885, and the National Institute on Aging (NIA) U01AG088407 (Hao Wang); as well as the University of Washington Environmental Health and Microbiome Research Center (EHMBRACE), an NIEHS P30ES030287 Environmental Health Sciences Center (EHSC) Pilot Award, and the Oregon State University College of Agricultural Sciences Environmental and Molecular Toxicology (CAS-EMT) Startup Fund (Julia Yue Cui). This work was also supported by a Global Grants for Gut Health Award from Nature and Yakult (Sean Gibbons).

